# Representation-aware calibration enables statistically reliable protein structure search

**DOI:** 10.64898/2026.11.03.764110

**Authors:** Lingfeng Shi, Han Zhou, Yifan Yang, Yang Young Lu

**Author notes:** These authors contributed equally to this work (Lingfeng Shi, Han Zhou).

## Abstract

Searching large databases of predicted protein structures can reveal remote homologs, but reliable discovery requires accurate significance estimates. We show that native E-values reported by protein structure search tools such as Foldseek and Reseek can be unreliable, leading to false discoveries or missed homologs. To address this problem, we introduce Struct-Caliper, which provides reliable query-specific p-values for protein structure search. The method uses representation-aware query randomization to generate background queries and compares observed alignment scores with the highest scores from randomized searches of the same database. In simulations, SG p-values generally kept reported false positive at or below the chosen cutoff under the evaluated randomization procedures. In searches of 15,177 ASTRAL40 structures, SG p-values increased Foldseek homolog recovery by 18.2% with slightly fewer false discoveries compared with native E-values at a cutoff of 0.01. For Reseek, false discoveries even decreased by 99.5%. Across 73 queries searched against more than two million AlphaFoldDB structures, SG p-values increased the mean recovery of DALI-supported matches by 23.5% at the same cutoff. These results show that Struct-Caliper can improve the reliability of structure searches and help prioritize discoveries for further biological analysis.

## 1 Introduction

Advances in protein structure prediction [1–3] have made three-dimensional structural models available for hundreds of millions of proteins [4]. Searching these models offers new opportunities to identify remote homologs and infer protein function, as structural similarity can persist even when sequence similarity is difficult to detect [5]. However, conventional methods such as DALI [6] and TM-align [7] compare three-dimensional protein structures directly, making exhaustive searches computationally expensive at this scale. Foldseek [8] and Reseek [9] address this challenge by transforming structure into a sequence of representations that encode residue-level structural features. Foldseek uses the 3Di structural alphabet, whereas Reseek combines multiple residue-level features into a larger alphabet. These representations allow alignment strategies for sequence search to be adapted to structural comparisons.

Finding similar structures efficiently, however, does not ensure that the significance reported for each match is reliable. To assess the significance of these sequence-like alignments, Foldseek and Reseek use E-values, following the same general framework as BLAST [10, 11]. An E-value estimates how many alignments scoring at least as highly as an observed match would occur by chance in a database search. Its reliability depends on whether the underlying statistical model accurately predicts how often such similarities arise by chance [12]. In our previous work, we showed that BLASTp E-values can either overestimate or underestimate significance, potentially making similarities between unrelated proteins appear significant or causing true homologs to be overlooked [13]. These findings highlight the need to validate significance estimates rather than assume that search methods provide naturally reliable statistical evidence. To address this problem in sequence search, we introduced Studentized-Gumbel (SG) p-values, which assess an observed alignment score against the highest scores obtained by searching randomized versions of the query against the same database [13]. The resulting p-values allow us to control the risk of false positives in database searches, improving the reliability of reported matches.

Extending SG p-values to structure search requires a suitable way to generate randomized queries. Amino-acid sequences can be shuffled while preserving their length and composition, but shuffling three-dimensional coordinates disrupts protein geometry. An alternative is to randomize the residue-level representations used by structure search methods. This randomization must disrupt patterns associated with homology while preserving the background properties needed to assess whether an alignment could arise by chance. Here we introduce Struct-Caliper, a framework that uses representation-aware query randomization to calibrate the statistical significance of protein structure searches (Fig. 1). For both Foldseek and Reseek, randomized queries are generated directly in the residue-level representations used for alignment, with the procedures described in Section 4.3. We then search the randomized representations against the same database and use the highest alignment scores to calculate SG p-values for the observed matches [13].

**Fig. 1:**
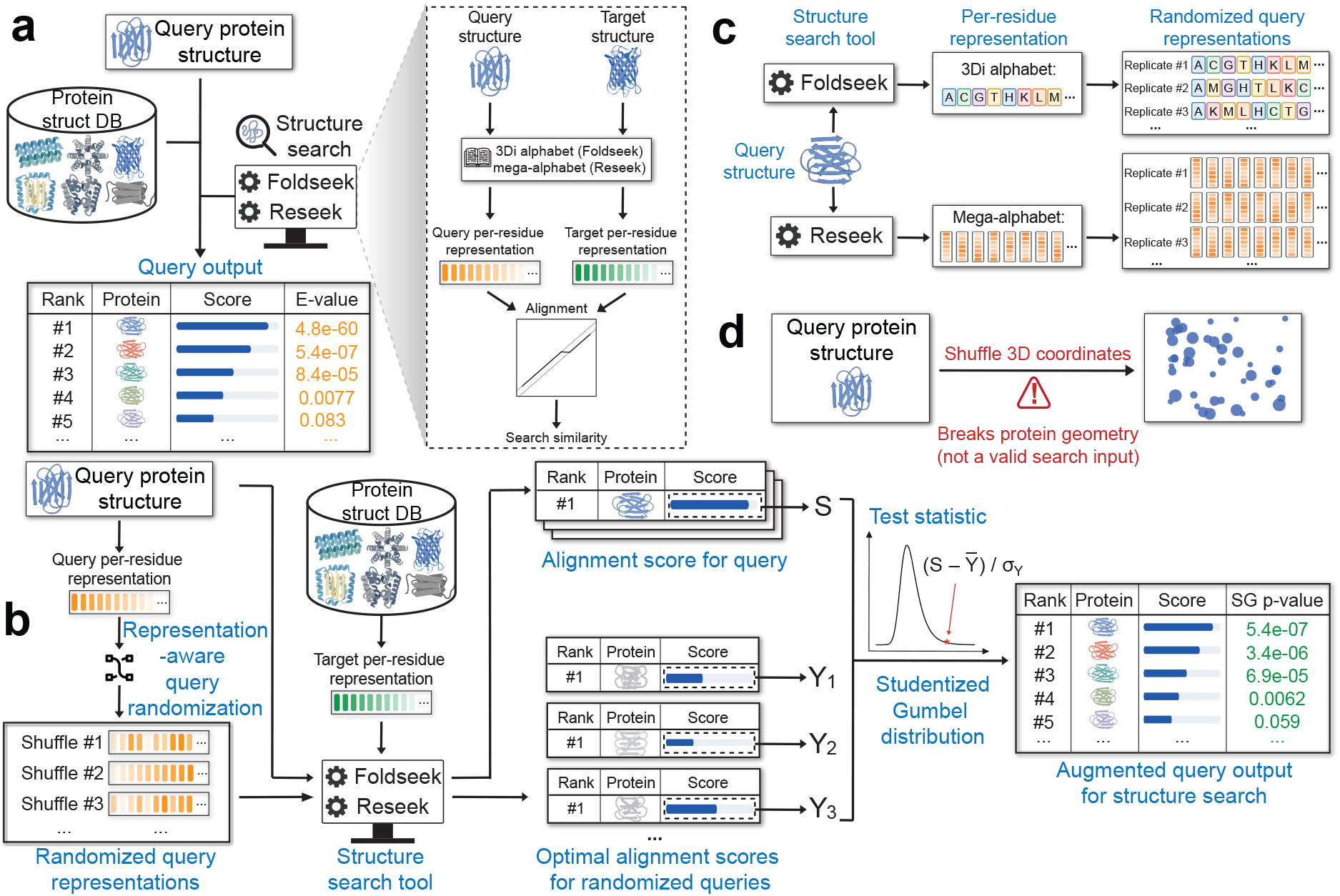
Overview of Struct-Caliper. **(a)** Foldseek and Reseek identify similar structures by aligning residue-level representations. **(b)** Struct-Caliper uses the highest scores from randomized-query searches against the same database to calculate Studentized-Gumbel p-values for observed matches. **(c)** Representation-aware query randomization operates on the amino-acid and 3Di sequences used by Fold-seek and the residue-level feature profiles used by Reseek. **(d)** Direct coordinate shuffling disrupts protein geometry and produces invalid search inputs.

We evaluate the reliability and practical benefits of Struct-Caliper through simulations and real applications. The simulations show that E-values reported by Foldseek and Reseek can overestimate or underestimate significance, whereas Struct-Caliper p-values control the risk of false positives at the specified significance levels. In homology searches across 15,177 ASTRAL40 structures, these p-values recover more homologs with substantially fewer false positives than E-values at the same significance levels. When used to pre-filter approximately 2 million AlphaFoldDB structures, Struct-Caliper reduces the number of candidates requiring detailed structural comparison while retaining DALI-supported matches. Together, these results demonstrate that Struct-Caliper improves the reliability of homology searches and prioritizes candidates for further analysis, supporting protein function annotation and discovery at scale.

## 2 Results

### 2.1 Simulations support the validity of Struct-Caliper SG p-values

We assessed whether Struct-Caliper SG p-values provide reliable false-positive control by comparing them with native E-values in randomized searches. For each query and search method, we performed 200,000 randomized searches against the same database of 15,177 ASTRAL40 structures [14]. The five query structures had lengths of 30, 87, 120, 151, and 288 residues, with details provided in Supplementary Section S2.1. Query representations were randomized as described in Section 4.3. We evaluated both Foldseek and Reseek, with Foldseek using its default mode combining 3Di and amino-acid information. Each SG p-value was calculated using *m* = 50 background scores following Section 4.4.

In these randomized searches, valid p-values should ensure that the probability of reporting any false discovery does not exceed the chosen cutoff [13]. We defined a discovery as a match whose E-value or p-value was at or below the cutoff threshold. Under the setting of randomized search, any such discovery represents a false positive. For example, a cutoff threshold of 0.01 should yield at least one discovery in no more than 1% of randomized searches on average. We therefore compared each cutoff with the fraction of searches reporting at least one discovery. Fractions consistently above the cutoff indicate inflated significance that may detect more false positives than expected, whereas substantially lower fractions indicate conservative behavior that may reduce sensitivity to true homologs. We applied this criterion to both Foldseek and Reseek native E-values and Struct-Caliper SG p-values (Fig. 2).

**Fig. 2:**
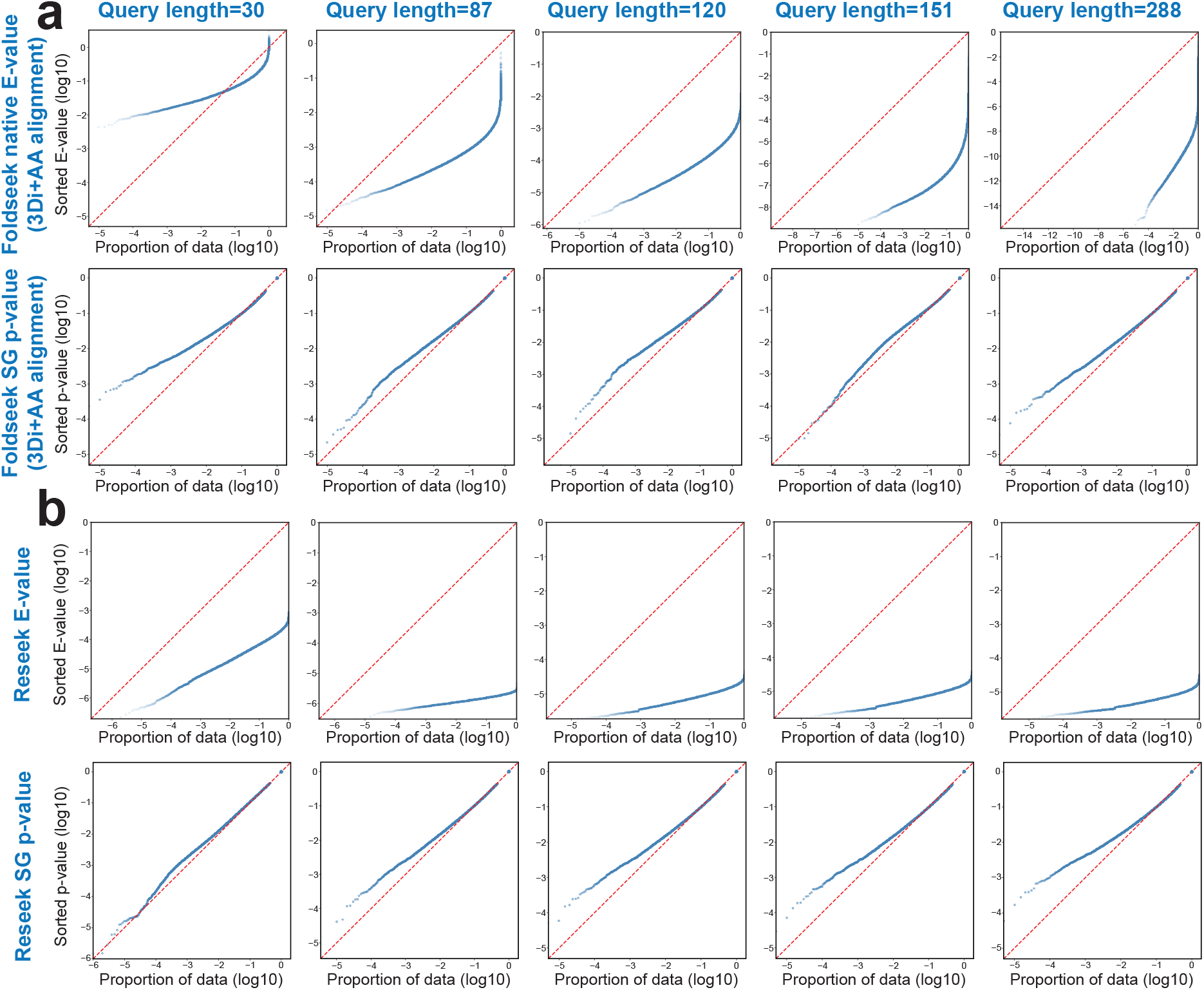
Native E-values can overestimate or underestimate the significance of structural search. **(a)** Native E-values and SG p-values for Foldseek. **(b)** Corresponding results for Reseek. For each method, the upper and lower rows show native E-values and Struct-Caliper SG p-values, respectively. Results are based on 200,000 randomized searches per query and search method against the ASTRAL40 database. Columns correspond to query lengths of 30, 87, 120, 151, and 288 residues from left to right. Sorted values are plotted against the cumulative proportion of randomized searches, with both axes on a log_10_ scale. The red dashed diagonal marks equality between reported values and empirical frequencies. Curves below the diagonal indicate overestimated significance and more false positives than expected, whereas curves above it indicate conservative estimates that may reduce sensitivity to true homologs.

Foldseek’s native E-values could either overestimate or underestimate significance, depending on the query (Fig. 2**a**). For the 30-residue query, stringent cutoffs yielded discoveries in a smaller fraction of searches than the cutoff would allow. Such conservative estimates may limit homolog discovery by requiring stronger alignment evidence than necessary. For the four longer queries, however, Foldseek reported false discoveries more often than expected across a wide range of E-value cutoffs. Significance inflation was particularly pronounced for the 151- and 288-residue queries. Thus, using the same E-value cutoff may lead to missed homologs for some queries and too many false discoveries for others. We observed similar problems when Foldseek used only 3Di structural information, without amino-acid information (Supplementary Fig. S1).

Reseek’s native E-values substantially overestimated significance across all five queries (Fig. 2**b**). At an E-value cutoff of 10^*−*3^, nearly all randomized searches would have reported at least one discovery. However, with correctly calibrated E-values, no more than 0.1% of randomized searches should report a discovery on average at this cutoff [13]. This discrepancy shows that even very stringent native E-values did not provide the expected protection against false discoveries. Significance inflation also persisted when we evaluated Reseek’s native p-values (Supplementary Fig. S2).

In contrast, Struct-Caliper SG p-values showed behavior consistent with validity across all five queries and both search methods (Fig. 2**a**,**b**). The fraction of randomized searches reporting at least one discovery was generally close to or below the chosen cutoff threshold. This behavior held even for queries whose native E-values most strongly overestimated significance. Although some settings remained conservative, valid p-values need only keep the false-positive probability at or below the cutoff [13]. Together, these simulations support the validity of Struct-Caliper SG p-values and their use for per-query false-positive control under the evaluated randomization procedures.

### 2.2 Struct-Caliper improves the reliability of homology discovery in ASTRAL40

We next evaluated whether Struct-Caliper SG p-values could improve homolog discovery and reduce false discoveries in searches of the ASTRAL40 database [14]. Each of the 15,177 structures was searched against the full database using Foldseek and Reseek. We compared discoveries reported at the same cutoff threshold of 0.01 for native E-values and SG p-values (Fig. 3). Pairs sharing a SCOP family or superfamily were classified as homologous, whereas pairs from different folds were classified as non-homologous and counted as false discoveries when reported.

**Fig. 3:**
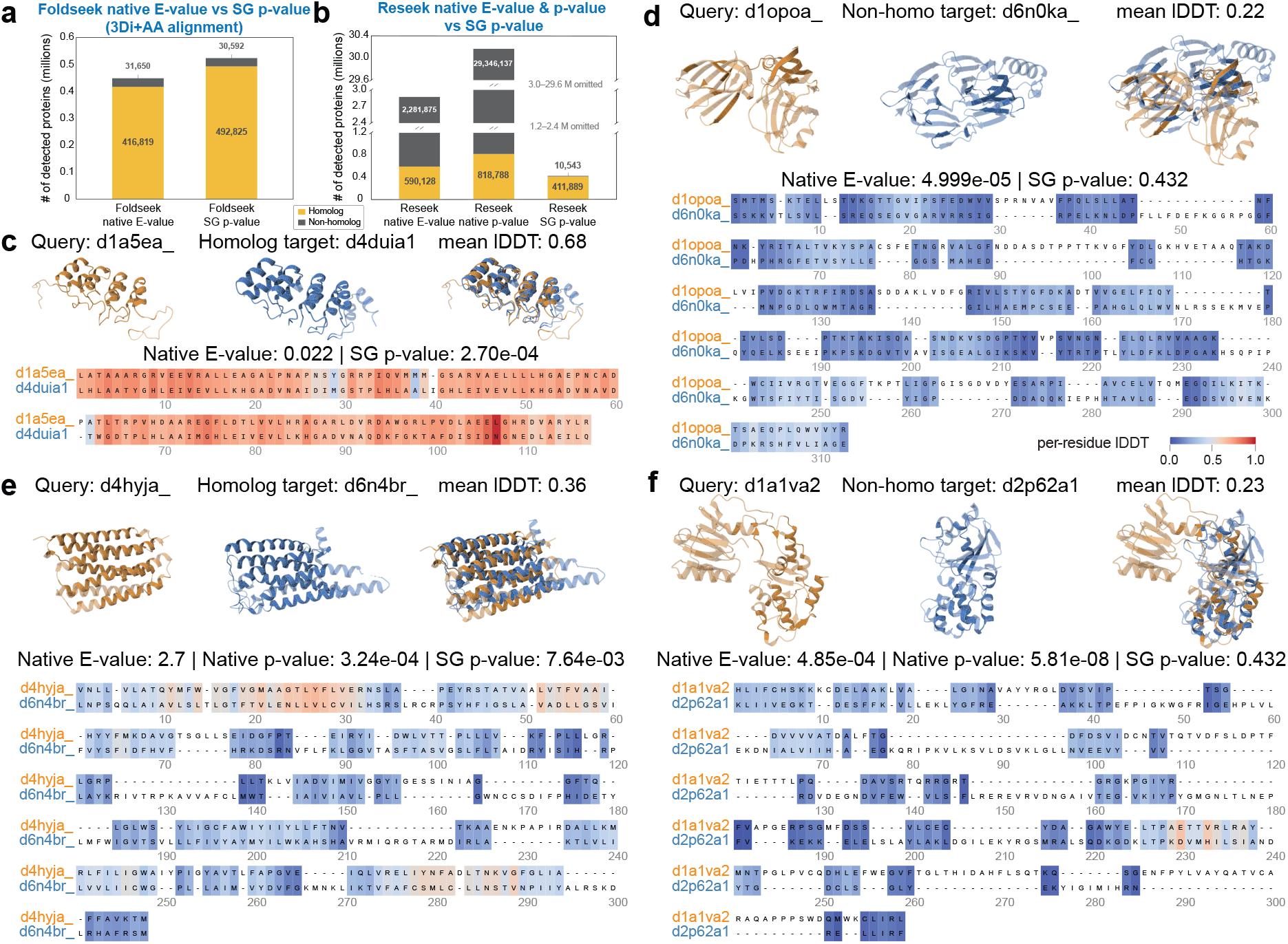
SG p-values increase homolog recovery for Foldseek and reduce false discoveries for Reseek. **(a)** Numbers of homologous and non-homologous discoveries reported using native E-values and Struct-Caliper SG p-values for Foldseek. **(b)** Corresponding results for Reseek, with native p-values also shown. Native E-values and SG p-values were thresholded at 0.01. Counts represent query–target pairs summed across all 15,177 ASTRAL40 queries. Pairs with uncertain homology labels were excluded. **(c**,**d)** 1. Examples from Foldseek’s 3Di-only alignment showing a homologous pair missed by native E-values and recovered using SG p-values, and a benchmark-negative pair reported using native E-values and excluded using SG p-values. **(e**,**f)** Corresponding examples for Reseek. Each example shows the query structure in orange, the target structure in blue, their superposition, and the sequence alignment returned by the search method. Mean lDDT values summarize local structural agreement, with higher values indicating better agreement.

For Foldseek, SG p-values increased homolog discovery while reducing false discoveries compared to native E-values. With the default alignment mode combining 3Di and amino-acid information, homologous discoveries increased from 416,819 using native E-values to 492,825 using SG p-values, an increase of 18.2% (Fig. 3**a**). False discoveries decreased slightly from 31,650 to 30,592. We also evaluated 3Di-only alignment to examine the benefits of significance calibration when amino-acid substitution scores are omitted from alignment scoring. In this mode, homologous discoveries increased from 384,473 to 443,333 an increase of 15.3%, while false discoveries decreased from 44,273 to 17,250, a reduction of 61.0% (Supplementary Fig. S3). The gain in homolog recovery is consistent with the simulations in Section 2.1, which showed that native E-values could be conservative for some queries.

To illustrate these effects at the level of individual matches, we selected examples from the 3Di-only analysis (Fig. 3**c**,**d**). These examples show how SG calibration can recover missed homologs and exclude benchmark-negative matches when alignment scoring uses structural representations alone, complementing the aggregate evaluation of Foldseek’s default 3Di+AA mode. The homologous pair `d1a5ea` and `d4duia1` showed local structural agreement with a mean lDDT of 0.68, but its native E-value of 0.022 exceeded the discovery cutoff (Fig. 3**c**). Its SG p-value of 2.70 *×* 10^*−*4^ allowed this pair to be reported. Conversely, the nonhomologous pair `d1opoa` and `d6n0ka` received a native E-value of 4.999 *×* 10^*−*5^ and would have been reported as a discovery (Fig. 3**d**). Its SG p-value of 0.432 excluded this false discovery. Additional examples are provided in Supplementary Fig S4 and Supplementary Fig S5.

For Reseek, the main benefit of SG p-values was a substantial reduction in false discoveries (Fig. 3**b**). False discoveries decreased from 2,281,875 using native E-values to 10,543 using SG p-values, a reduction of 99.5%. The additional comparison using Reseek’s native p-values yielded an even larger number of false discoveries, totaling 29,346,137. These results are consistent with the significance overestimation observed for native Reseek E-values and p-values in randomized searches (Section 2.1 and Supplementary Fig. S2). Despite loss of some homologous discoveries, SG p-values recovered individual Reseek homologs that native E-values missed. The homologous pair `d4hyja` and `d6n4br` received a native E-value of 2.7 and was therefore excluded (Fig. 3**e**). Its native p-value of 3.24 *×* 10^*−*4^ and SG p-value of 7.64 *×* 10^*−*3^ both allowed the pair to be reported. In contrast, the non-homologous pair `d1a1va2` and `d2p62a1` received a native E-value of 4.85 *×* 10^*−*4^ and a native p-value of 5.81 *×* 10^*−*8^, both of which would have led to a false discovery (Fig. 3**f**). Its SG p-value of 0.432 excluded this false discovery. Additional examples are provided in Supplementary Fig S6 and Supplementary Fig S7. These examples show that SG p-values can recover missed homologs while reducing non-homologous discoveries substantially.

### 2.3 Struct-Caliper improves sensitivity in large-scale AlphaFoldDB searches

We next evaluated whether the gains observed in ASTRAL40 extended to searches against millions of predicted protein structures. We used 73 SCOPe protein structures as queries and a clustered subset of AlphaFoldDB from a published benchmark as the target database [15]. The target database contained more than two million predicted structures, over 100 times the size of ASTRAL40. We focused on Foldseek because Reseek searches were computationally impractical at this scale. Each query was searched using Foldseek’s default mode combining 3Di and amino-acid information. We compared discoveries reported using native E-values and Struct-Caliper SG p-values from the same search results.

Unlike ASTRAL40, the AlphaFoldDB benchmark lacked homology labels, so we used DALI scores to assess structural support for reported discoveries [15]. DALI compares three-dimensional protein structures directly, with higher scores indicating greater structural similarity [6]. Following the benchmark, we considered a pair structurally supported when its DALI score met a threshold chosen separately for each query. The threshold was selected using the Kneedle algorithm, which identifies the bend in the query’s ranked DALI scores where their decline begins to slow [16]. These reference labels allowed us to assess recovery for queries with both few and many structurally supported targets (Fig. 4**a**,**b**).

**Fig. 4:**
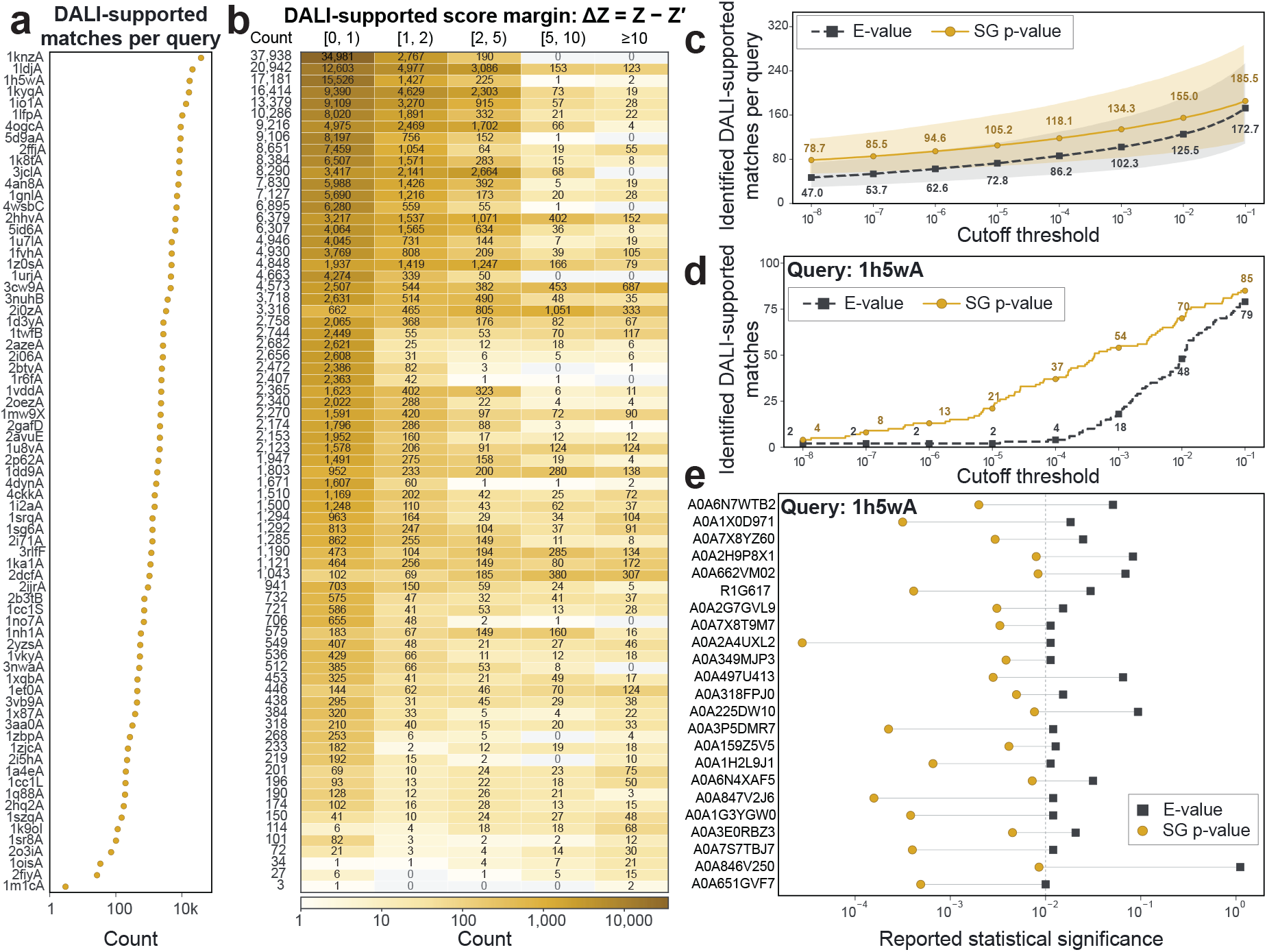
SG p-values recover more structurally supported matches in large-scale AlphaFoldDB searches. We evaluated 73 protein queries against more than two million predicted structures. Matches were considered structurally supported when their DALI scores met a reference threshold chosen separately for each query. **(a)** Number of DALI-supported matches among the evaluated candidates for each query. **(b)** These matches grouped by how far their DALI scores exceeded the query-specific threshold. **(c)** Mean number of DALI-supported matches recovered per query at different cutoffs. Shaded regions show 95% confidence intervals at each cutoff. **(d)** Corresponding results for query `1h5wA`. **(e)** The 23 DALI-supported matches for this query recovered using SG p-values but excluded using native E-values at a cutoff threshold of 0.01. Connected points show the two values for each match, and the vertical dashed line marks the cutoff.

SG p-values increased the recovery of DALI-supported matches across most queries in this larger database. At a cutoff threshold of 0.01, the mean number of supported discoveries per query increased from 125.5 using native E-values to 155.0 using SG p-values, an increase of 23.5% (Fig. 4**c**). The higher average recovery persisted across the evaluated cutoffs. Query `1h5wA` illustrated how SG p-values recovered structurally supported matches that native E-values missed. SG p-values reported 70 supported discoveries, compared with 48 using native E-values (Fig. 4**d**). SG p-values recovered 23 supported matches excluded by native E-values, whereas one supported match was reported only using native E-values (Fig. 4**e**). Additional examples are provided in Supplementary Fig. S8. These results extend the gains in recovery observed in ASTRAL40 to searches against millions of predicted protein structures.

## 3 Discussion

We introduced Struct-Caliper to improve the reliability of statistical significance estimates in protein structure search. The method uses representation-aware query randomization to construct query-specific backgrounds and evaluates observed alignment scores against the highest scores from randomized searches of the same database. This extends the SG p-value framework developed for sequence search to structural comparisons without requiring randomized three-dimensional coordinates [13]. In simulations, native Foldseek and Reseek E-values could overestimate or underestimate significance, whereas SG p-values generally kept the reported false positives at or below the chosen cutoff. In ASTRAL40, SG p-values increased Foldseek homolog recovery by 18.2% with slightly fewer false discoveries and reduced Reseek false discoveries by 99.5%, although some homologous discoveries were also excluded. The gain in Foldseek recovery extended to searches against more than two million AlphaFoldDB structures, where the mean number of DALI-supported discoveries increased by 23.5%.

The reliability of Struct-Caliper depends on whether the randomized queries provide an appropriate background for the search [13]. The current procedures preserve selected properties of each representation while disrupting dependencies associated with homology. Choosing which properties to preserve is consequential, because removing too much local structure may underestimate chance similarities, whereas preserving patterns associated with homology may make the resulting p-values unnecessarily conservative. Our simulations support validity under the evaluated procedures, while broader validation across protein architectures and search settings remains necessary.

Several extensions could broaden the applicability of Struct-Caliper. First, alternative randomizations could help determine which local dependencies are important for reliable significance estimation. Algorithms such as uShuffle, which preserve the counts of short sequence patterns exactly, provide a way to investigate this question for structural alphabets [17]. Second, integrating the randomized queries into batched GPU searches could reduce the computational cost of constructing a separate background for each query, building on recent acceleration of MMseqs2 and Foldseek [18]. Finally, studies involving many queries would benefit from extending error control beyond individual searches. Procedures for controlling the false discovery rate under dependence provide a foundation for assessing discoveries across related queries and shared target databases [19, 20].

Our results demonstrate that changing the significance estimate can alter which relationships are reported even when the underlying alignments remain the same. Reliable significance assessment therefore has a direct role in determining which candidates receive further biological attention. By helping recover overlooked homologs and reduce false discoveries, Struct-Caliper can support the prioritization of candidates for evolutionary analysis, functional annotation, and experimental validation. The framework also provides a basis for evaluating statistical reliability as new structural representations and search methods are developed.

## 4 Methods

### 4.1 Protein structure search methods

Protein structure search compares a query structure against a structure database to identify structurally similar proteins. We evaluated Foldseek [8] and Reseek v2.8 [9], which encode protein structures as ordered, residue-level representations. Both methods use these representations to identify structural matches through sequence-style alignment but differ in how they encode and score structural information.

Foldseek [8] encodes tertiary interactions using the 20-state 3Di structural alphabet. For each residue, it extracts ten geometric features describing its relationship with a spatial neighbor, including relative orientation, spatial distance, and sequence separation. A neural network trained to capture evolutionarily conserved structural information maps these features to a structural representation. Each protein is thus represented by a 3Di sequence alongside its amino-acid sequence. Foldseek uses MMseqs2 [21] to identify candidate targets and then applies Smith-Waterman local alignment. We evaluated both 3Di-only and combined 3Di and amino-acid alignment, an E-value reporting cutoff of 10^9^, and a maximum of 1,000 reported hits per query. Original queries retained the native scoring settings. For randomized queries, reverse-score subtraction and compositional-bias correction were disabled. For all queries, we used floating-point bit scores calculated from Foldseek’s predicted Gumbel parameters.

Reseek [9] represents each residue using amino-acid identity and eight structural features describing backbone conformation and relationships with spatial neighbors. Each structural feature is assigned to one of 16 states, yielding a combined mega-alphabet of approximately 10^11^ possible states. Rather than estimating a substitution score for every pair of mega-alphabet states, Reseek learns a separate scoring matrix for each feature from reference structural alignments. Scores for aligned residues are weighted sums of these feature-specific scores. Local alignments are obtained using Smith-Waterman or a BLAST-like procedure with gap penalties. Reseek also calculates an alignment quality score which combines the alignment score with local structural agreement and corrections for protein length and reversed-query similarity. All searches used the most sensitive search mode, with the same alignment and scoring parameters for original and randomized queries. We used alignment quality scores to calculate p-values.

### 4.2 Calculation of native E-values

An E-value estimates how many alignments scoring at least as highly as an observed match would occur by chance in a database search [13]. In the classical BLAST framework, the database-wide E-value for an alignment score *s* is approximately:

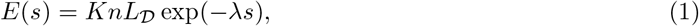

where *n* is the query length, *L*_*D*_ is the total number of residues in the target database, and *K* and *λ* are parameters that characterize the background score distribution for the chosen scoring system [12, 13].

Foldseek and Reseek calculate their native E-values using different background models [8, 9]. Foldseek uses a neural network to predict the parameters of its background score distribution from the query’s amino-acid composition and length [8]. Reseek instead converts alignment quality scores to E-values using an empirical score distribution [9]. Further details are provided in Supplementary Section S1.1. In our analyses, we used the E-values reported for the original queries by each structure search method.

### 4.4 Representation-aware query randomization

Struct-Caliper constructs query-specific background distributions using representation-aware query randomization. These randomizations are designed to disrupt patterns associated with homology while preserving relevant background properties of each representation. Let **x** = (*x*_1_, …, *x*_*n*_) denote an ordered query representation, where *x*_*i*_ is the representation at residue position *i* and *n* is the query length.

For Foldseek, **x** denotes either the amino-acid sequence or the 3Di sequence from 20-state 3Di structural alphabet. We fitted separate first-order Markov models to these sequences, estimating transition probabilities from the frequencies of adjacent pairs. Each randomized sequence began with the first state of the corresponding original sequence, and subsequent states were sampled according to the fitted transition probabilities for the preceding state. The amino-acid and 3Di sequences were generated independently using their respective models. This approach captures dependencies between adjacent positions while disrupting longer-range sequence patterns. For Foldseek, this procedure explicitly models adjacent-state dependencies that uniform positional shuffling does not preserve.

For Reseek, **x** denotes the complete residue-level feature profiles across all residue positions. For each replicate, we sampled a uniform random permutation *π* of the residue positions and reordered the profiles as 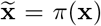. This permutation changes only the order of the profiles, preserving each residue-level representation and the relationships among its features while disrupting dependencies between residues along the sequence. We modified Reseek to internally generate randomized queries directly during searches, avoiding the overhead of creating a separate database on disk.

### 4.4 Calculation of SG p-values and family-wise error rate control

Struct-Caliper assesses the statistical significance of structural alignment scores using query-specific background distributions. For each query, we generated *m* = 50 randomized queries using the representation-aware query randomization procedure described in Section 4.3 and searched them against the same target database. We recorded the highest score from each randomized search to obtain the background scores *Y*_1_, …, *Y*_*m*_. Let *S*_*i*_ denote the observed score between the original query and target *i*. We studentized each score as:

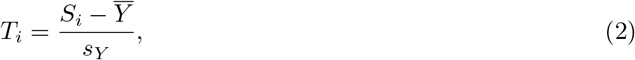

where 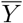 and *s*_*Y*_ are the sample mean and sample standard deviation of the background scores.

Assuming the background scores are independent draws from a common Gumbel distribution, we assessed *T*_*i*_ using the Studentized-Gumbel (SG) reference distribution [13]. This reference depends only on *m* and accounts for uncertainty in the background mean and standard deviation estimated from the randomized searches. We calculated the p-value for each observed match as

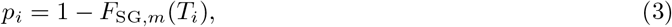

where *F*_SG,*m*_ is the cumulative distribution function of the SG distribution. We used the precomputed reference distribution for *m* = 50, obtained by Monte Carlo simulation with importance sampling [13]. Further details about SG distribution are provided in Supplementary Section S1.3.

Because each background score is the highest score from a randomized search against the entire database, the resulting p-values account for comparisons across all targets. When these p-values are valid, reporting matches with *p*_*i*_ ≤ *α* controls the family-wise error rate at level *α* [13]. This means that, for each query, the probability of reporting even one false positive is at most *α*. More details are provided in Supplementary Section S1.5. We evaluated the validity of these p-values using the randomized-search experiments described in Section 2.1.

## 5 Data Availability

This study used the ASTRAL40 dataset [14] and the AlphaFoldDB benchmark described by **(author?)** [15]. Processed data, including query identifiers, reference labels, background scores from randomized searches, and search results with alignment scores, native E-values, and Struct-Caliper SG p-values, are available through Zenodo at https://doi.org/10.5281/zenodo.23872787.

## 6 Code Availability

Source code for representation-aware query randomization, SG p-value calculation, and the analyses and figure generation reported in this study is available at https://github.com/batmen-lab/Struct-Caliper. The repository includes the modified Reseek implementation, scripts for running Foldseek searches, and instructions for reproducing the experiments.

## 7 Acknowledgements

This work was supported by startup funding from the University of Wisconsin–Madison RISE AI Initiative, a Google Research Award and the NVIDIA Academic Grant Program.

## 8 Author contributions

L.S. implemented the code, set up and preprocessed the datasets, and performed the analysis. H.Z. implemented the code for the homology discovery analysis and performed the analysis. L.S., H.Z. and Y.Y.L. prepared the figures and participated in the discussion. Y.Y.L. wrote the manuscript. All authors participated the discussion. All authors reviewed the manuscript.

