## Supplement for "Representation-aware calibration enables statistically reliable protein structure search"

### S1 Method details

#### S1.1 Calculation of native E-values

An E-value estimates the expected number of alignments that would occur by chance with scores at least as high as an observed match [4]. In the classical BLAST framework, the database-wide E-value for an alignment score $s$ is approximately:

$$E(s) = KnL_{\mathcal{D}} \exp(-\lambda s), \tag{1}$$

where $n$ is the query length, $L_{\mathcal{D}}$ is the total number of residues in the target database, and $K$ and $\lambda$ describe the background score distribution for the chosen scoring system [2, 4]. Foldseek and Reseek use different background models to calculate their native E-values [3, 1].

**Foldseek E-values.** Foldseek models background alignment scores using a query-dependent Gumbel distribution [3]. Its scoring procedure applies compositional-bias correction and subtracts the reversed-query alignment score from the forward score. A neural network predicts the Gumbel parameters from query composition and length. The network is trained using 100,000 structures sampled from AlphaFoldDB. Training targets are obtained by maximum-likelihood fitting of Gumbel distributions to alignment scores against shuffled database sequences. The network contains two fully connected hidden layers with 32 units each and two output units for the distribution parameters. The predicted parameters provide an initial E-value estimate, denoted $E_0$. Calibration using scrambled-sequence searches identifies a log-linear relationship between false positives per query and predicted E-values. The estimated slope is 0.32 and Foldseek incorporates this empirical adjustment through the power transformation:

$$E_{\text{Foldseek}} = E_0^{0.32}. \tag{2}$$

The reported E-value therefore combines query-dependent Gumbel parameter prediction with an empirical calibration correction [3].

**Reseek E-values.** Reseek combines the local alignment score and geometric agreement into an alignment statistic [1]. This statistic is defined as:

$$t = \delta\Delta + \frac{\alpha s - \beta s_{\text{rev}}}{L + \lambda}, \tag{3}$$

where $s$ is the local alignment score and $\Delta$ is lDDT-mu, a measure of local structural agreement. The quantity $L$ denotes the mean query and target length. The score $s_{\text{rev}}$ provides a reversed-query background correction. The offset $\lambda$ moderates length normalization for short proteins. The parameters $\alpha$, $\beta$, $\delta$, and $\lambda$ belong to Reseek's scoring model and are distinct from the BLAST parameters in Eq. (1). The alignment quality score is obtained through the transformation

$$\text{AQ} = \left[1 + \frac{1}{2} 10^{(a+bt)/10}\right]^{-1}. \tag{4}$$

The scoring and transformation parameters are estimated using SCOP40 training data and assessed by held-out validation. AQ ranges from zero to one, with larger values indicating stronger alignment evidence. Reseek estimates native p-values using an empirical piecewise log-linear calibration of the alignment statistic $t$. In Reseek v2.8, the E-value is calculated as

$$E_{\text{Reseek}}(t) = N_{\text{ref}}\, p_{\text{Reseek}}(t), \tag{5}$$

where $p_{\text{Reseek}}(t)$ is the native p-value for an alignment with statistic $t$, and $N_{\text{ref}}$ is the number of structures in the reference database used for E-value scaling. The Reseek v2.8 implementation fixes $N_{\text{ref}}$ at the SCOP40c reference-database size of 8,340 structures, without adjusting this factor to the number of targets in the searched database.

#### S1.2 Studentization removes location and scale parameters

The SG reference distribution can be reused across queries because studentization removes the location and scale parameters of the assumed null distribution [4]. Here, we give the argument for database-wide maximum alignment scores. For a given query and target database, let $S$ denote a maximum score generated under the null and let $Y_1, \ldots, Y_m$ denote the maxima from $m$ independently randomized queries. Assume that $S, Y_1, \ldots, Y_m$ are independent draws from the same location-scale family with location $\mu$ and scale $\beta > 0$. For $m \geq 2$, define

$$\bar{Y} = \frac{1}{m}\sum_{j=1}^{m} Y_j, \qquad \sigma_Y = \left[\frac{1}{m-1}\sum_{j=1}^{m}(Y_j - \bar{Y})^2\right]^{1/2}. \tag{6}$$

When $\sigma_Y > 0$, the studentized score is $T = (S - \bar{Y})/\sigma_Y$.

Each draw from a location-scale family can be written as a shifted and rescaled draw from a common reference distribution. We can therefore write

$$S = \mu + \beta Z_0, \qquad Y_j = \mu + \beta Z_j, \qquad j = 1, \ldots, m, \tag{7}$$

where $Z_0, Z_1, \ldots, Z_m$ are independent draws from a distribution that does not depend on $\mu$ or $\beta$. Let $\bar{Z}$ and $\sigma_Z$ denote the sample mean and standard deviation of $Z_1, \ldots, Z_m$, using the same definitions as above. It follows that $\bar{Y} = \mu + \beta\bar{Z}$ and $\sigma_Y = \beta\sigma_Z$. Substituting these expressions gives

$$T = \frac{S - \bar{Y}}{\sigma_Y} = \frac{\beta(Z_0 - \bar{Z})}{\beta\sigma_Z} = \frac{Z_0 - \bar{Z}}{\sigma_Z}. \tag{8}$$

The location parameter cancels when the background mean is subtracted, and the scale parameter cancels when the result is divided by the background standard deviation. Thus, the distribution of $T$ depends on the common distributional shape and the sample size $m$, but not on $\mu$ or $\beta$.

When the common distribution is Gumbel, the resulting reference distribution is $\mathrm{SG}_m$ [4]. A single table of its tail probabilities can therefore be used for queries with different null locations and scales, provided that the same $m$ and standard-deviation convention are used. This property allows Foldseek and Reseek to share the $\mathrm{SG}_{50}$ table while using separate query-specific background samples. It does not remove differences in distributional shape, so applying the SG reference still requires an appropriate randomized background and an adequate Gumbel approximation for the maximum scores.

#### S1.3 Computation of the Studentized-Gumbel distribution

Struct-Caliper uses the Studentized-Gumbel (SG) distribution introduced in our previous work to assess alignment scores relative to a query-specific background [4]. Here, we describe the reference distribution and the numerical procedure used to construct the published SG lookup table.

**Definition of the reference distribution.** For a given query, let $Y_1, \ldots, Y_m$ denote the highest alignment scores obtained by searching $m$ independently randomized queries against the same target database under the same search settings. Their sample mean and standard deviation are

$$\bar{Y} = \frac{1}{m}\sum_{j=1}^{m} Y_j, \qquad \sigma_Y = \left[\frac{1}{m-1}\sum_{j=1}^{m}(Y_j - \bar{Y})^2\right]^{1/2}. \tag{9}$$

The SG model assumes that a database-wide maximum score $S$ obtained under the null and the background scores $Y_1, \ldots, Y_m$ are independent draws from the same Gumbel distribution. When $\sigma_Y > 0$, the studentized score is

$$T = \frac{S - \bar{Y}}{\sigma_Y}. \tag{10}$$

Subtracting the sample mean removes the location parameter, and dividing by the sample standard deviation removes the scale parameter. The distribution of $T$ therefore depends only on $m$ and is denoted by $\mathrm{SG}_m$ [4]. Its upper-tail probability is

$$G_m(t) = \Pr(T \geq t). \tag{11}$$

This reference distribution accounts for variation in both the null maximum score and the background mean and standard deviation estimated from a finite sample. We use $m = 50$ in Struct-Caliper and therefore evaluate scores using the $\mathrm{SG}_{50}$ reference table.

**Numerical estimation of tail probabilities.** Direct simulation is inefficient for estimating very small tail probabilities because large values of $T$ occur rarely. The published $\mathrm{SG}_{50}$ table was therefore computed using importance sampling, which generates large score values more frequently and corrects for this change through numerical weights [4]. The calculation used $N = 10^{10}$ independent simulation replicates. In each replicate $r$, a sample of $m = 50$ background values was drawn from a Gumbel distribution with location 0 and scale 3, giving a sample mean $\bar{Y}_r$ and standard deviation $\sigma_{Y,r}$. An independent score $S_r$ was drawn from a broader Gumbel distribution with location 33 and scale 15 to improve coverage of the upper tail.

Writing the Gumbel density with location $\mu$ and scale $\beta > 0$ as

$$g_{\mu,\beta}(s) = \frac{1}{\beta} \exp\left[-\frac{s-\mu}{\beta} - \exp\left(-\frac{s-\mu}{\beta}\right)\right], \tag{12}$$

each replicate produces a studentized score and an importance weight

$$T_r = \frac{S_r - \bar{Y}_r}{\sigma_{Y,r}}, \qquad w_r = \frac{g_{0,3}(S_r)}{g_{33,15}(S_r)}. \tag{13}$$

The density ratio corrects for drawing $S_r$ from a different distribution than the background sample. The upper-tail probability is then estimated by

$$\widehat{G}_m(t) = \frac{1}{N} \sum_{r=1}^{N} w_r \mathbf{1}\{T_r \geq t\}, \tag{14}$$

where the indicator is one when $T_r \geq t$ and zero otherwise. Thus, the weights of simulated scores at or above $t$ are summed and divided by the total number of replicates. This computation uses simulated Gumbel values rather than protein database searches and is performed once for each choice of $m$.

**Lookup table and alignment p-values.** The published calculation accumulated simulated values in bins of width 0.001 over the interval from 0 to 45. Following its numerical precision assessment, the final lookup table was restricted to values up to 34, below the reported precision boundary of 34.599 [4]. The estimated tail probability at this cutoff is $\widehat{G}_{50}(34) \approx 8.716 \times 10^{-15}$.

For an observed alignment score $S_i$ between the original query and target $i$, we calculate

$$t_i = \frac{S_i - \bar{Y}}{\sigma_Y} \tag{15}$$

and obtain the p-value from the reference table using

$$p_i = \begin{cases} 1, & t_i \leq 0, \\ \widehat{G}_{50}(t_i), & 0 < t_i \leq 34, \\ \widehat{G}_{50}(34), & t_i > 34. \end{cases} \tag{16}$$

Within the tabulated interval, the lookup uses the grid value at or immediately below $t_i$. Scores at or below the background mean are assigned a p-value of one, whereas scores above the upper cutoff receive the smallest tabulated p-value rather than an extrapolated estimate [4]. The same reference table can be used for both Foldseek and Reseek because the SG distribution depends only on $m$ under the common-Gumbel model. Its application still requires an appropriate randomized background and an adequate Gumbel approximation for the database-wide maximum scores.

#### S1.4 Algorithm for computing SG p-values in structure search

Algorithm 1 adapts our SG p-value procedure to representation-based structure search [4]. For a query $q$ and target database $\mathcal{D}$, let $S_1, \ldots, S_k$ denote the observed alignment scores. The algorithm uses representation-aware query randomization to generate $m$ randomized query representations, searches each against $\mathcal{D}$, and records its highest score. The resulting background scores $Y_1, \ldots, Y_m$ are used to studentize every observed score from the original query.

For Foldseek, randomized amino-acid and 3Di sequences are generated independently from separate first-order Markov models fitted to the query. For Reseek, complete residue-level feature profiles are shuffled among positions using our modified search implementation. We use Foldseek bit scores or Reseek alignment quality scores, following the method-specific scoring procedures described in the main Methods. The target database remains unchanged throughout the procedure.

Let $\widehat{G}_m(u)$ denote the tabulated upper-tail probability of the $\mathrm{SG}_m$ distribution at grid value $u$. The table must correspond to the number of randomized queries and the standard-deviation convention used in the algorithm. In our analyses, $m = 50$, the grid spacing is $h = 0.001$, and the largest tabulated value is $t_{\max} = 34$, with $\widehat{G}_{50}(34) \approx 8.716 \times 10^{-15}$ [4]. For a studentized score within this range, the algorithm uses the grid value at or immediately below it. Scores at or below zero receive a p-value of one, and scores above $t_{\max}$ receive the smallest tabulated p-value rather than an extrapolated estimate. The calculation requires a finite maximum score from each randomized search and a positive background standard deviation. Construction of the reference table is described in Section S1.3.

**Algorithm 1** SG p-values for representation-based structure search

**Input** Query $q$, target database $\mathcal{D}$, observed scores $S_1, \ldots, S_k$
**Input** Chosen search method and its representation-aware query randomization procedure
**Input** Number of randomized queries $m \geq 2$ and matching table $\widehat{G}_m$
**Input** Table spacing $h$ and upper limit $t_{\max}$
**Output** SG p-values $p_1, \ldots, p_k$

**for** $j = 1, \ldots, m$ **do**
    Generate a randomized representation $\widetilde{q}_j$ from $q$ using the specified procedure
    Search $\widetilde{q}_j$ against $\mathcal{D}$ using the chosen method
    $Y_j \leftarrow$ the highest score from that search
**end for**
$\bar{Y} \leftarrow \frac{1}{m} \sum_{j=1}^{m} Y_j$
$\sigma_Y \leftarrow \left[ \frac{1}{m-1} \sum_{j=1}^{m} (Y_j - \bar{Y})^2 \right]^{1/2}$
Proceed only when all $Y_j$ are finite and $\sigma_Y > 0$
**for** $i = 1, \ldots, k$ **do**
    $t_i \leftarrow (S_i - \bar{Y})/\sigma_Y$
    **if** $t_i \leq 0$ **then**
        $p_i \leftarrow 1$
    **else if** $t_i > t_{\max}$ **then**
        $p_i \leftarrow \widehat{G}_m(t_{\max})$
    **else**
        $u_i \leftarrow h\lfloor t_i/h \rfloor$
        $p_i \leftarrow \widehat{G}_m(u_i)$
    **end if**
**end for**
**return** $(p_1, \ldots, p_k)$

#### S1.5 Control of the family-wise error rate

Struct-Caliper uses SG p-values to select significant structural alignments while accounting for the multiple matches considered in a query's database search. Following the argument introduced in our previous work

[4], we show how a valid p-value for the highest null alignment score provides control of the family-wise error rate (FWER). Here, FWER is the probability of reporting at least one false positive under the specified null model.

**Selection rule.** For a single query, let $S_1, \ldots, S_k$ denote the observed alignment scores against the target database. Let $Y_1, \ldots, Y_m$ denote the highest scores obtained from $m$ independently randomized queries searched against the same database under the same settings, with $m = 50$ in our analyses. Write $\bar{Y}$ and $\sigma_Y$ for their sample mean and standard deviation, with $\sigma_Y > 0$. For the theoretical argument, let $G_m$ denote the exact upper-tail probability function of the $\mathrm{SG}_m$ distribution. We assign each alignment the p-value

$$p_i = G_m\left(\frac{S_i - \bar{Y}}{\sigma_Y}\right) \tag{17}$$

and report it as significant when $p_i \leq \alpha$, where $0 < \alpha < 1$ is the chosen significance threshold. Numerical evaluation of these p-values is described in Section S1.3.

**Error-control argument.** Let $\mathcal{I}_0$ denote the indices of alignments for which the null hypothesis is true. When this set is nonempty, define the highest null score and its corresponding p-value as

$$S_0 = \max_{i \in \mathcal{I}_0} S_i, \qquad p_0 = G_m\left(\frac{S_0 - \bar{Y}}{\sigma_Y}\right). \tag{18}$$

Assume that $p_0$ is valid, meaning that $\Pr(p_0 \leq \alpha) \leq \alpha$ for every $0 < \alpha < 1$. Because $G_m$ decreases as the score increases and all alignments use the same background mean and standard deviation, the highest null score has the smallest p-value among null alignments. Thus, $p_0 = \min_{i \in \mathcal{I}_0} p_i$, and

$$\mathrm{FWER} = \Pr\left(\min_{i \in \mathcal{I}_0} p_i \leq \alpha\right) = \Pr(p_0 \leq \alpha) \leq \alpha. \tag{19}$$

If there are no null alignments, the FWER is zero. Under the stated validity condition, no additional correction for the number of matches within this search is needed because the reference distribution already accounts for the highest score across the database. The argument does not require independence among alignment scores for different targets.

**Assumptions and scope.** Under the complete null, where every alignment is null, $S_0$ is the highest score across the entire database. If $S_0$ and $Y_1, \ldots, Y_m$ are independent draws from the same Gumbel distribution, then $(S_0 - \bar{Y})/\sigma_Y$ follows the $\mathrm{SG}_m$ distribution and $p_0$ is valid [4]. The error-control argument also holds when $p_0$ is conservative rather than exactly uniform. When true and null matches coexist, the same guarantee requires the randomized background to remain a valid or conservative reference for the highest null score $S_0$. Validity under the complete null alone does not establish this additional property. In practice, the guarantee also depends on the suitability of the randomized background and the accuracy of the numerical SG tail probabilities. Finally, this guarantee applies to one query searched against a specified database. Controlling the probability of any false positive across multiple queries requires an additional adjustment across queries.

### S2 Result details

#### S2.1 Proteins used for simulation study

The five query structures used in the simulation study were selected from ASTRAL40 (SCOPe version 2.08) to span sequence lengths from 30 to 288 residues. Table S1 lists their domain identifiers, database annotations, and full amino acid sequences.

Table S1: Protein domains used as queries in the simulation study.

<table>
<tr><th>Domain ID<br>SCOPe code</th><th>Length<br>(residues)</th><th>Protein / domain<br>Source organism</th><th>PDB ID<br>Chain:residues</th></tr>
<tr><td>d1zavu1<br>a.108.1.1</td><td>30</td><td>Ribosomal protein L7/12, oligomerisation (N-terminal) domain<br>Thermotoga maritima</td><td>1ZAV<br>U:1–30</td></tr>
<tr><td colspan="4">MTIDEIIEAI EKLTVSELAE LVKKLEDKFG</td></tr>
<tr><td>d1stza1<br>a.4.5.51</td><td>87</td><td>Heat-inducible transcription repressor HrcA, N-terminal domain<br>Thermotoga maritima</td><td>1STZ<br>A:14–100</td></tr>
<tr><td colspan="4">KLNDRQRKVL YCIVREYIEN KKPVSSQRVL EVSNIEFSSA TIRNDMKKLE YLGYIYQPHT SAGRIPTDKG LRFYYEEMLK<br>ISKETSE</td></tr>
<tr><td>d2ao9a1<br>a.4.1.17</td><td>120</td><td>Phage protein BC1890<br>Bacillus cereus</td><td>2AO9<br>A:13–132</td></tr>
<tr><td colspan="4">MMAKLDELKQ KLTAKQIQAA YLLVENELME SNNEEKRTQD EMANELGINR TTLWEWRTKN QDFIAFKSEV ADSFLAEKRE<br>QVYSKLMQLI LGPQPSVKAM QLYMQRFGLL TDKKVIEGDL</td></tr>
<tr><td>d1ku9a_<br>a.4.5.36</td><td>151</td><td>DNA-binding protein Mj223<br>Methanococcus jannaschii</td><td>1KU9<br>A:</td></tr>
<tr><td colspan="4">MIIMEEAKKL IIELFSELAK IHGLNKSVGA VYAILYLSDK PLTISDIMEE LKISKGNVSM SLKKLEELGF VRKVWIKGER<br>KNYYEAVDGF SSIKDIAKRK HDLIAKTYED LKKLEEKCNE EEKEFIKQKI KGIERMKKIS EKILEALNDL D</td></tr>
<tr><td>d2uy2a_<br>c.1.8.0</td><td>288</td><td>Automated matches<br>Saccharomyces cerevisiae</td><td>2UY2<br>A:</td></tr>
<tr><td colspan="4">ANTNIAVYWG QNSAGTQESL ATYCESSDAD IFLLSFLNQF PTLGLNFANA CSDTFSDGLL HCTQIAEDIE TCQSLGKKVL<br>LSLGGASGSY LFSDDSQAET FAQTLWDTFG EGTGASERPF DSAVVDGFDF DIENNNEVGY SALATKLRTL FAEGTKQYYL<br>SAAPQCPYPD ASVGDLLENA DIDFAFIQFY NNYCSVSGQF NWDTWLTYAQ TVSPNKNIKL FLGLPGSASA AGSGYISDTS<br>LLESTIADIA SSSSFGGIAL WDASQAFSNE LNGEPYVEIL KNLLTSAS</td></tr>
</table>

*Notes.* Metadata and sequences were extracted from the ASTRAL40 SCOPe 2.08 SEQRES file `astral-scopedom-seqres-gd-sel-gs-bib-40-2.08.fa.txt`. The SCOPe classification code is shown below each domain identifier; PDB denotes the Protein Data Bank. Chain and residue annotations follow the FASTA headers; `A:` indicates that no explicit residue range is given. Each metadata row is followed by the complete sequence in one-letter amino acid notation, grouped in blocks of ten residues (up to 80 residues per line). For `d2uy2a_`, "automated matches" is the description provided in the source database.

#### S2.2 More results of simulation

The additional simulations supported the validity of Struct-Caliper SG p-values for both Foldseek alignment modes and further showed that Reseek's native p-values overestimated significance. We followed the simulation protocol described in the main text, using 200,000 randomized searches per query and search configuration against the same database of 15,177 ASTRAL40 structures. The five queries had lengths of 30, 87, 120, 151, and 288 residues (Table S1), and each SG p-value was calculated using $m = 50$ background scores. As in the main analysis, we compared reported values with the fraction of randomized searches yielding at least one discovery at the corresponding cutoff.

Foldseek's native E-values showed query-dependent miscalibration in both 3Di-only alignment and combined 3Di and amino-acid alignment (Fig. S1**a,b**). For the 30-residue query, native E-values were conservative at stringent cutoffs. For the four longer queries, they overestimated significance across a wide range of cutoffs, with particularly pronounced deviations for the 151- and 288-residue queries. Thus, using only structural information did not resolve the miscalibration observed with combined 3Di and amino-acid alignment. In both modes, Struct-Caliper SG p-values generally followed or lay above the diagonal across all five queries, consistent with false-positive probabilities at or below the chosen cutoffs under the evaluated randomization procedures.

Reseek's native p-values also substantially overestimated significance across all five queries (Fig. S2, middle row). Both native E-values and native p-values lay well below the diagonal, indicating that small reported values occurred much more frequently than their corresponding cutoffs would allow. In contrast, Struct-Caliper SG p-values generally lay on or above the diagonal, with conservative behavior particularly apparent at stringent cutoffs for the four longer queries (Fig. S2, lower row). These additional comparisons support the use of Struct-Caliper SG p-values for per-query false-positive control under the evaluated simulation settings.

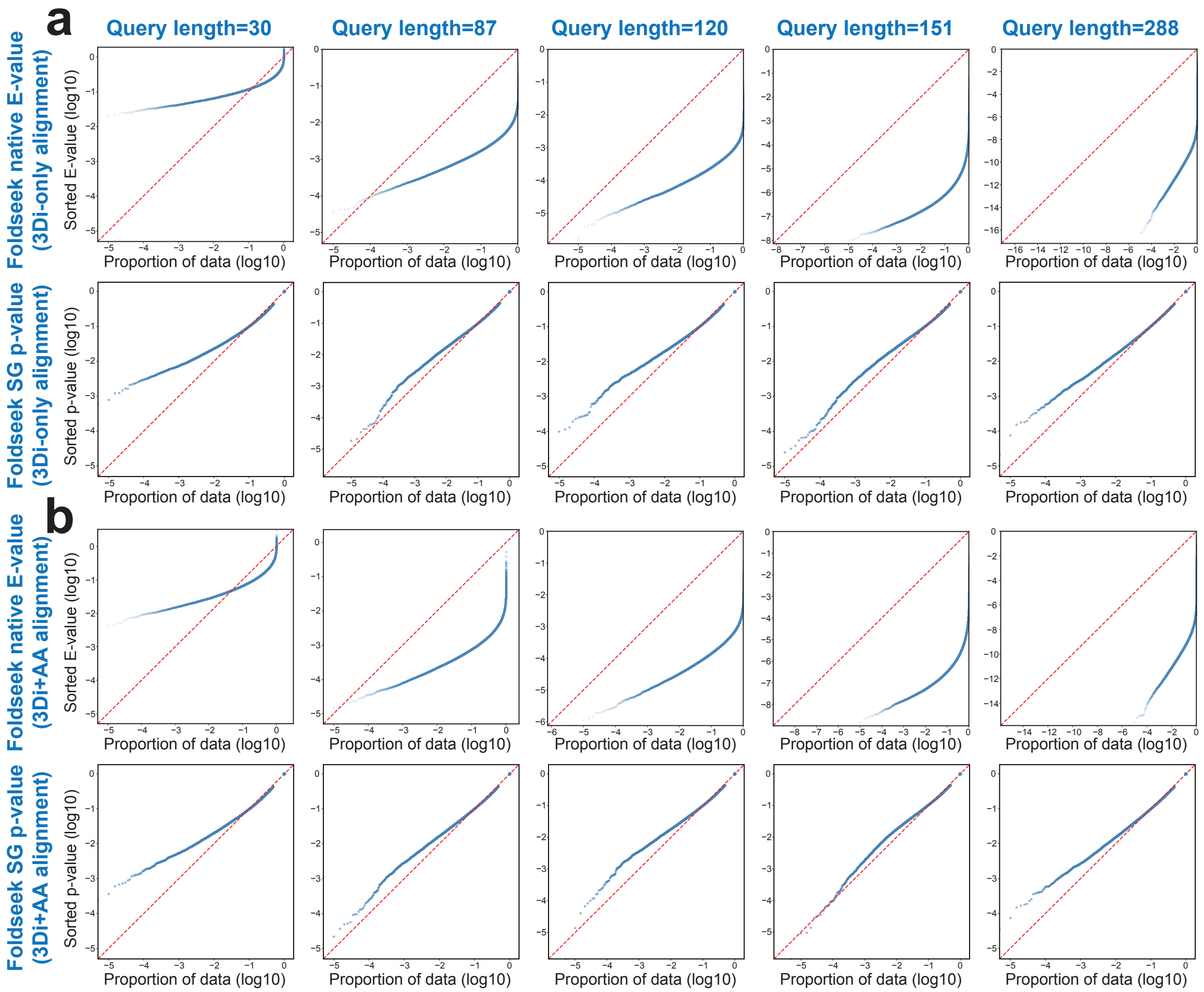

Figure S1: **Native E-values can overestimate or underestimate significance in both Foldseek alignment modes.** **(a)** Native E-values and Struct-Caliper SG p-values for Foldseek using only 3Di structural information. **(b)** Corresponding results for combined 3Di and amino-acid (AA) alignment. For each mode, the upper and lower rows show native E-values and Struct-Caliper SG p-values, respectively. Results are based on 200,000 randomized searches per query and alignment mode against the ASTRAL40 database of 15,177 structures, with $m = 50$ background scores used to calculate each SG p-value. Columns correspond to query lengths of 30, 87, 120, 151, and 288 residues from left to right. Sorted values on the vertical axis are plotted against the cumulative proportion of randomized searches on the horizontal axis, with both axes on a $\log_{10}$ scale. The red dashed diagonal marks equality between reported values and empirical frequencies. Curves below the diagonal indicate overestimated significance and more false positives than expected, whereas curves above it indicate conservative estimates that may reduce sensitivity to true homologs.

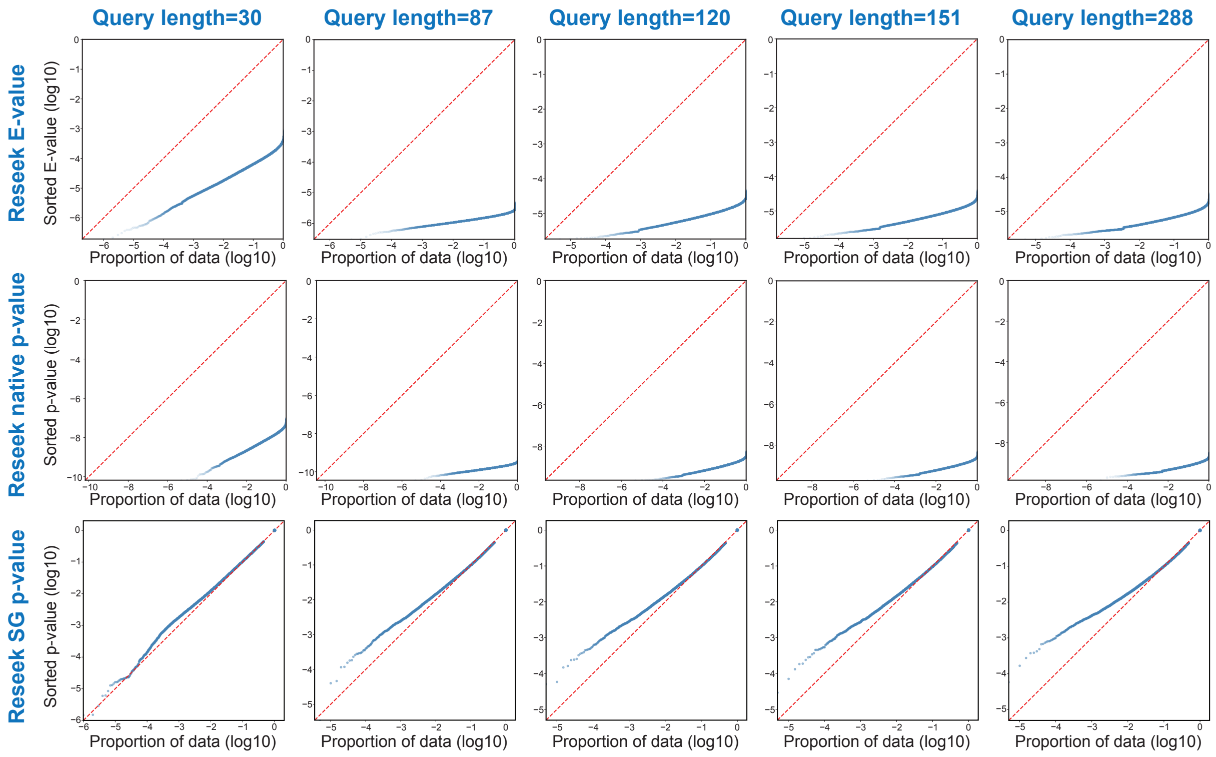

Figure S2: **Reseek's native E-values and p-values overestimate significance in randomized searches.** The upper, middle, and lower rows show Reseek native E-values, Reseek native p-values, and Struct-Caliper SG p-values, respectively. Results are based on 200,000 randomized searches per query against the ASTRAL40 database of 15,177 structures, with $m = 50$ background scores used to calculate each SG p-value. Columns correspond to query lengths of 30, 87, 120, 151, and 288 residues from left to right. Sorted E-values or p-values on the vertical axis are plotted against the cumulative proportion of randomized searches on the horizontal axis, with both axes on a $\log_{10}$ scale. The red dashed diagonal marks equality between reported values and empirical frequencies. Curves below the diagonal indicate overestimated significance and more false positives than expected, whereas curves above it indicate conservative estimates that may reduce sensitivity to true homologs.

#### S2.3 More results of ASTRAL40 search

The expanded ASTRAL40 analysis compared homolog recovery and false discoveries for both Foldseek alignment modes and for Reseek (Fig. S3). As in the main analysis, each of the 15,177 structures was searched against the full database; native E-values, Reseek native p-values, and Struct-Caliper SG p-values were all thresholded at 0.01. Pairs sharing a SCOP family or superfamily were classified as homologous, whereas pairs from different folds were classified as non-homologous. Pairs with uncertain homology labels were excluded from the counts.

For Foldseek 3Di-only alignment, SG p-values increased homologous discoveries from 384,473 to 443,333 (15.3%) while reducing non-homologous discoveries from 44,273 to 17,250 (61.0%; Fig. S3**a**, left). The corresponding comparison for combined 3Di and amino-acid alignment is shown alongside it (Fig. S3**a**, right). For Reseek, SG p-values yielded 411,889 homologous discoveries, compared with 590,128 using native E-values and 818,788 using native p-values. The respective numbers of non-homologous discoveries were 10,543, 2,281,875, and 29,346,137 (Fig. S3**b**). Thus, the reduction in Reseek false discoveries was accompanied by a reduction in homolog recovery.

Additional Foldseek examples showed how SG p-values recovered missed homologs and excluded non-homologous matches (Figs. S4 and S5). The homologous pair `d1914a1` and `d1914a2` had a mean lDDT of 0.66 but a native E-value of 0.012, just above the discovery cutoff. Its SG p-value of $2.22 \times 10^{-7}$ allowed the pair to be reported. Conversely, the non-homologous pair `d1opoa_` and `d6n0ka_` had a mean lDDT of 0.22 and a native E-value of $4.999 \times 10^{-5}$, whereas its SG p-value of 0.432 excluded the match.

Reseek examples showed that SG p-values could recover individual homologs despite the lower total homolog count (Fig. S6). For example, `d1nsca_` and `d1w0pa3` had a native E-value of 0.1575 but an SG p-value of $8.70 \times 10^{-3}$. All three recovered Reseek examples were also reported using native p-values at the 0.01 cutoff. In contrast, the three non-homologous examples in Fig. S7 had both native E-values and native p-values below 0.01, but SG p-values of 0.432. Their mean lDDT values ranged from 0.14 to 0.23, indicating limited local structural agreement.

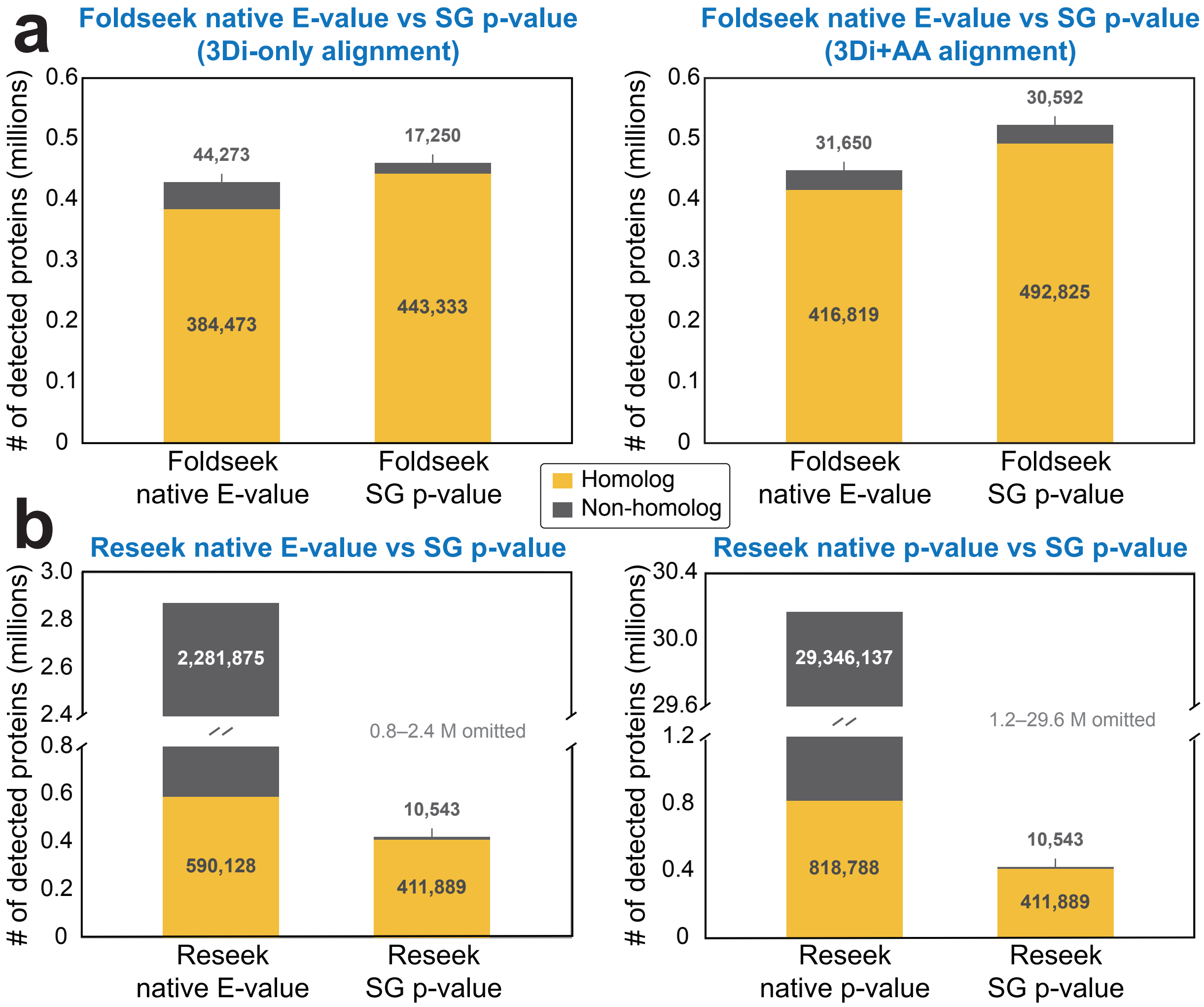

Figure S3: **Expanded comparison of homolog recovery and false discoveries in ASTRAL40. (a)** Numbers of homologous and non-homologous discoveries reported using native E-values and Struct-Caliper SG p-values for Foldseek with 3Di-only alignment (left) and combined 3Di and amino-acid alignment (right). **(b)** Corresponding Reseek comparisons using native E-values (left) and native p-values (right); the same SG p-value results are shown in both panels. Native E-values, Reseek native p-values, and SG p-values were all thresholded at 0.01. Counts represent query–target pairs summed across all 15,177 ASTRAL40 queries searched against the full database. Pairs sharing a SCOP family or superfamily were classified as homologous (yellow), and pairs from different folds as non-homologous (gray). Pairs with uncertain homology labels were excluded. Bar labels give exact counts, and vertical axes show counts in millions. The broken axes in **(b)** omit the indicated ranges.

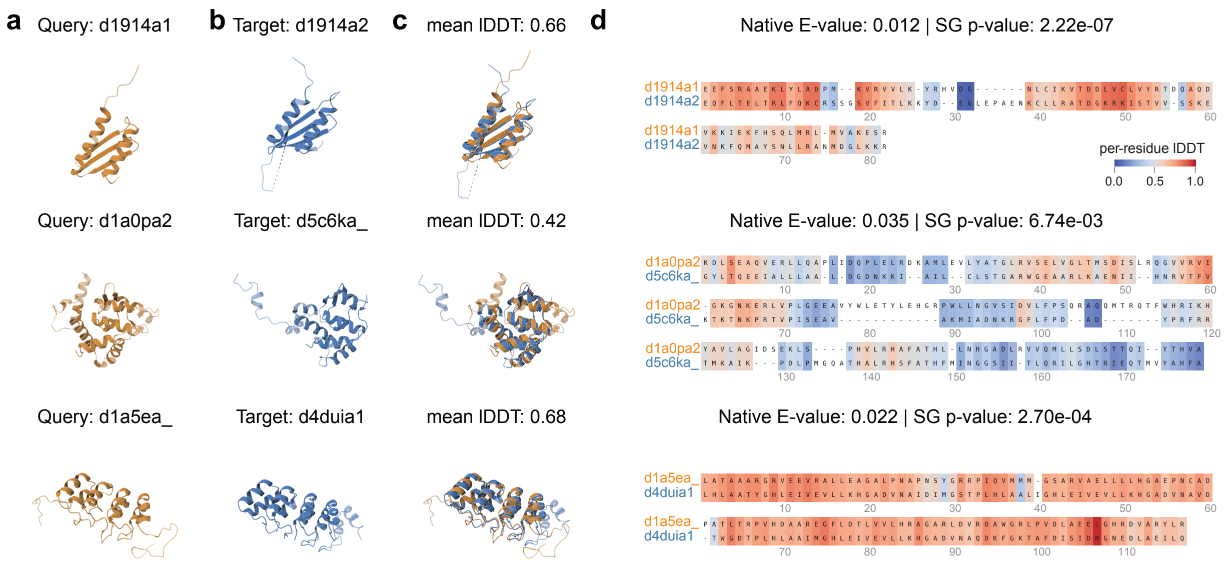

Figure S4: **Struct-Caliper recovers homologous pairs missed by Foldseek native E-values.** Each query and target share a SCOP family. All three pairs had native E-values above 0.01 but SG p-values below 0.01. Rows show `d1914a1`–`d1914a2`, `d1a0pa2`–`d5c6ka_`, and `d1a5ea_`–`d4duia1`, from top to bottom. Columns show **(a)** the query structure in orange, **(b)** the target structure in blue, **(c)** their superposition with the mean lDDT, and **(d)** the sequence alignment returned by Foldseek with the native E-value and Struct-Caliper SG p-value. Alignment shading indicates per-residue lDDT on the scale shown, from blue (0) to red (1). Mean lDDT values summarize local structural agreement, with higher values indicating better agreement.

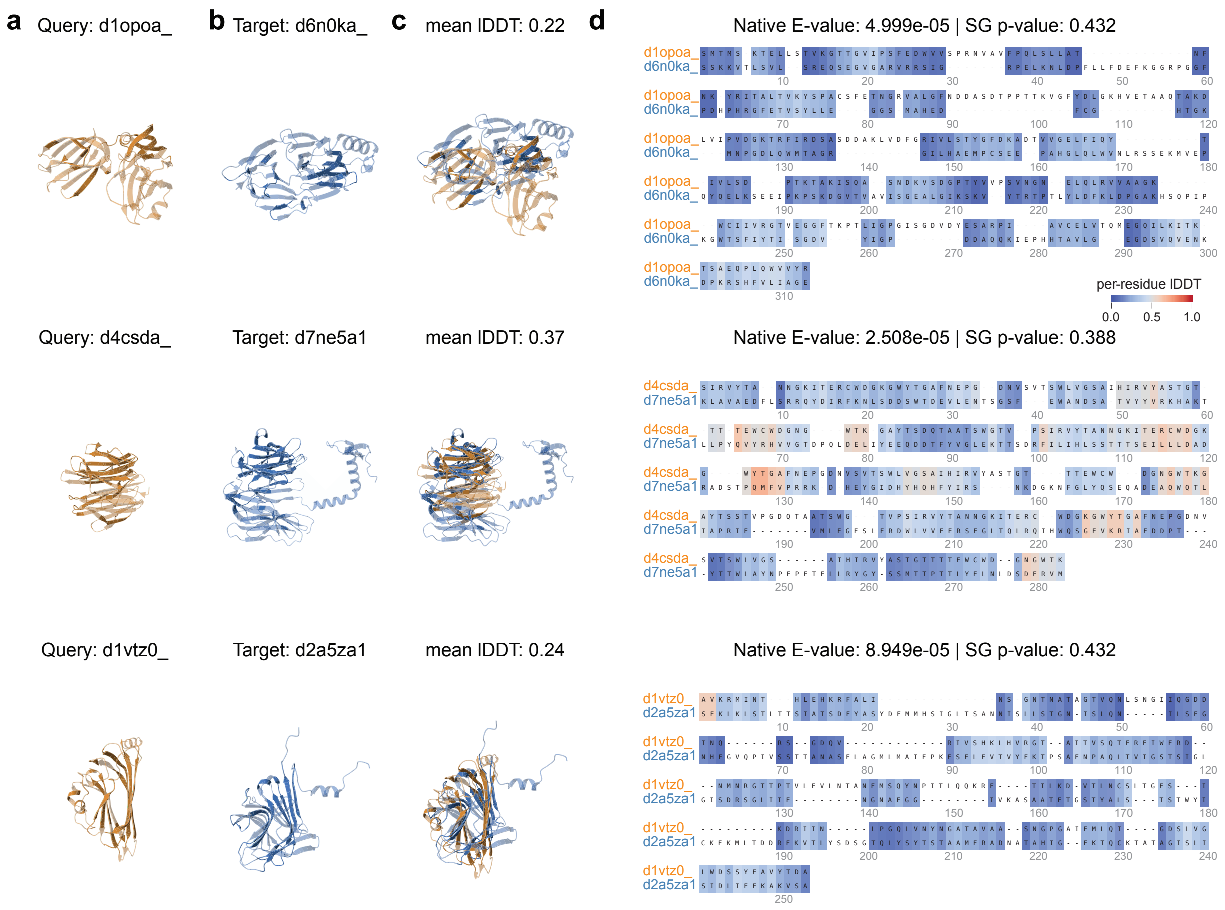

Figure S5: **Struct-Caliper excludes non-homologous pairs reported using Foldseek native E-values.** Each query and target belong to different SCOP folds. All three pairs had native E-values below 0.01 but SG p-values above 0.01. Rows show `d1opoa_–d6n0ka_`, `d4csda_–d7ne5a1`, and `d1vtz0_–d2a5za1`, from top to bottom. Columns show **(a)** the query structure in orange, **(b)** the target structure in blue, **(c)** their superposition with the mean lDDT, and **(d)** the sequence alignment returned by Foldseek with the native E-value and Struct-Caliper SG p-value. Alignment shading indicates per-residue lDDT on the scale shown, from blue (0) to red (1). Mean lDDT values summarize local structural agreement, with higher values indicating better agreement.

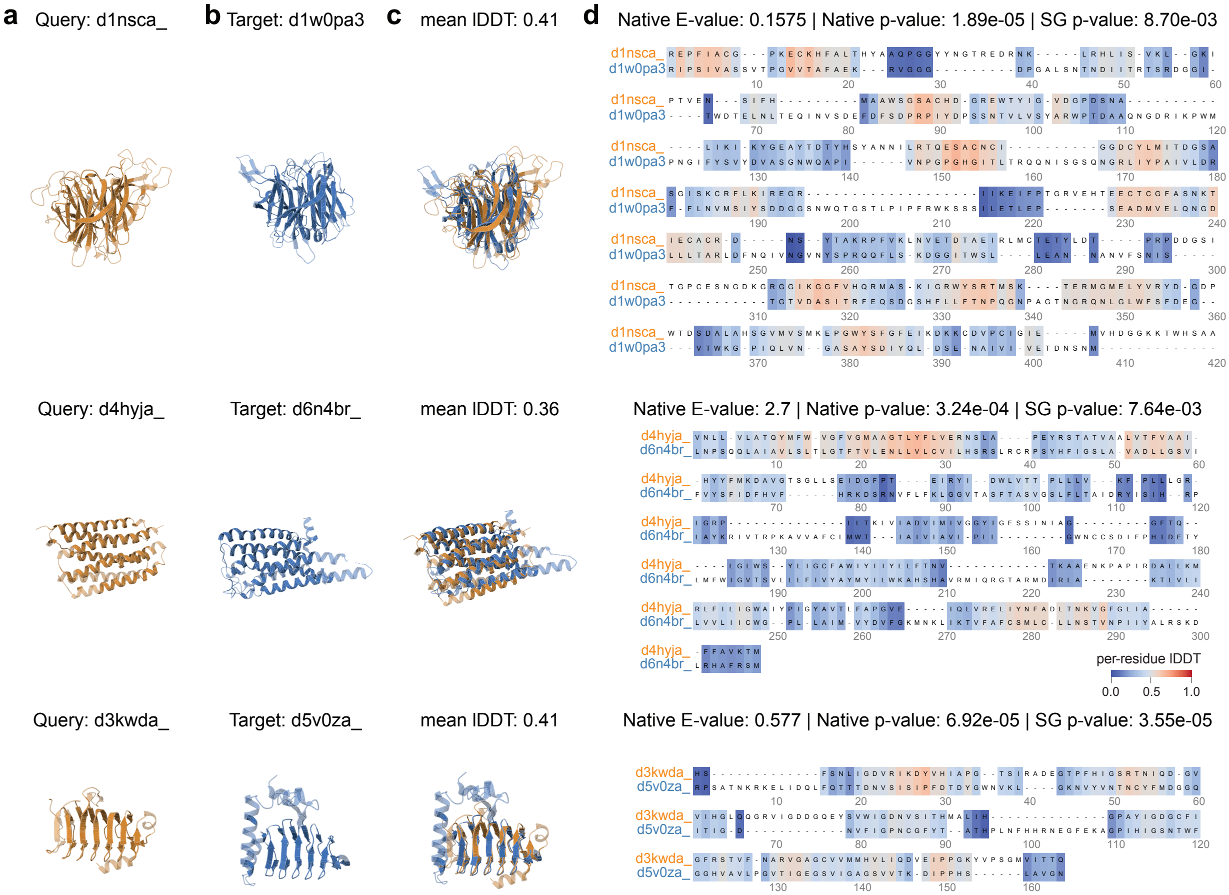

Figure S6: **Struct-Caliper recovers homologous pairs missed by Reseek native E-values.** Each query and target share a SCOP family. All three pairs had native E-values above 0.01 but SG p-values below 0.01; their native p-values were also below 0.01. Rows show `d1nsca_–d1w0pa3`, `d4hyja_–d6n4br_`, and `d3kwda_–d5v0za_`, from top to bottom. Columns show **(a)** the query structure in orange, **(b)** the target structure in blue, **(c)** their superposition with the mean lDDT, and **(d)** the sequence alignment returned by Reseek with the native E-value, native p-value, and Struct-Caliper SG p-value. Alignment shading indicates per-residue lDDT on the scale shown, from blue (0) to red (1). Mean lDDT values summarize local structural agreement, with higher values indicating better agreement.

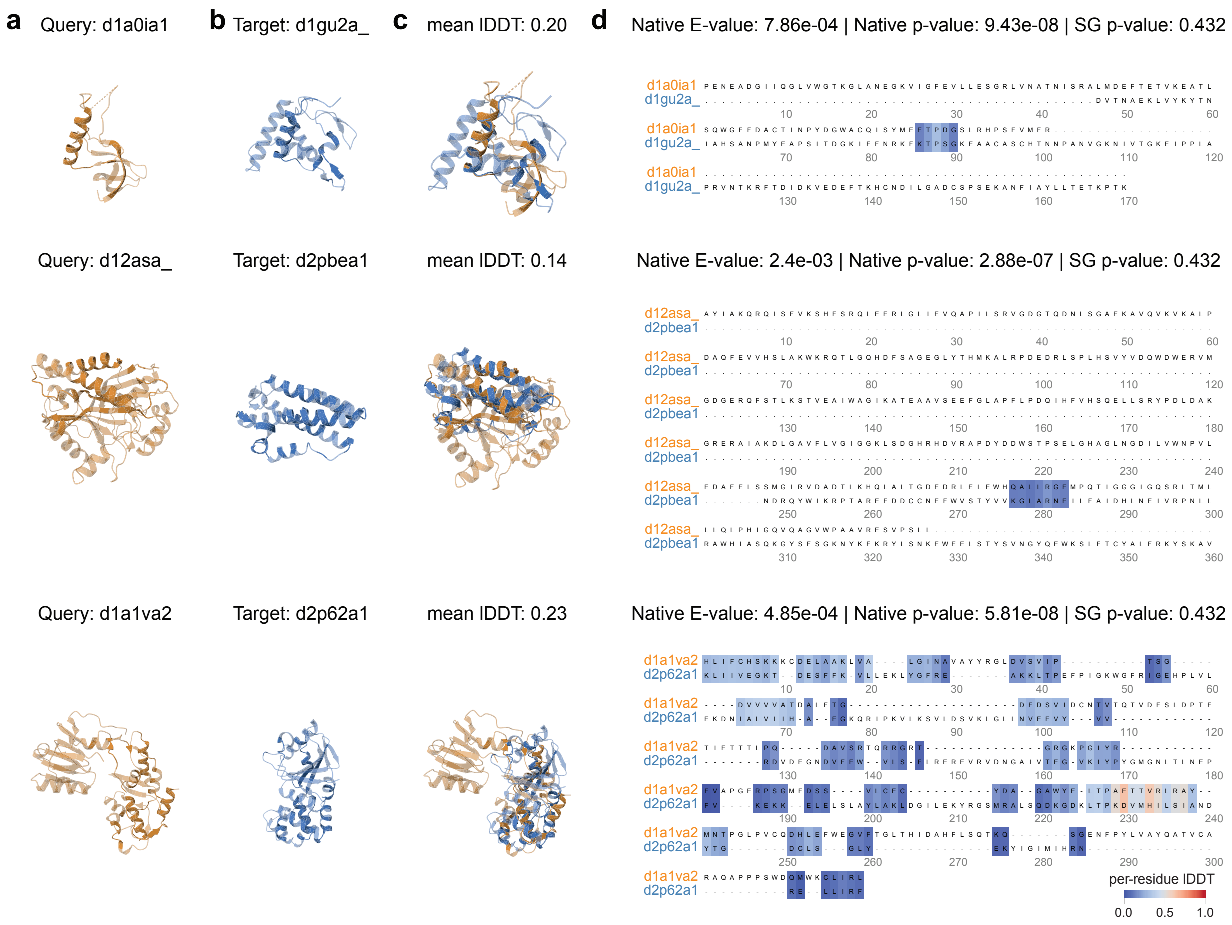

Figure S7: **Struct-Caliper excludes non-homologous pairs reported using Reseek native E-values and p-values.** Each query and target belong to different SCOP folds. All three pairs had native E-values and native p-values below 0.01 but SG p-values above 0.01. Rows show `d1a0ia1`–`d1gu2a_`, `d12asa_`–`d2pbea1`, and `d1a1va2`–`d2p62a1`, from top to bottom. Columns show **(a)** the query structure in orange, **(b)** the target structure in blue, **(c)** their superposition with the mean lDDT, and **(d)** the sequence alignment returned by Reseek with the native E-value, native p-value, and Struct-Caliper SG p-value. Alignment shading indicates per-residue lDDT on the scale shown, from blue (0) to red (1). Mean lDDT values summarize local structural agreement, with higher values indicating better agreement.

#### S2.4 More results of AlphaFoldDB search

To examine how the recovery gains varied among individual AlphaFoldDB queries, we expanded the query-level comparison shown in the main text to 15 examples (Fig. S8). All searches used Foldseek's default mode combining 3Di and amino-acid information against the same database of more than two million predicted structures. DALI support was defined using the same query-specific reference thresholds as in the main analysis.

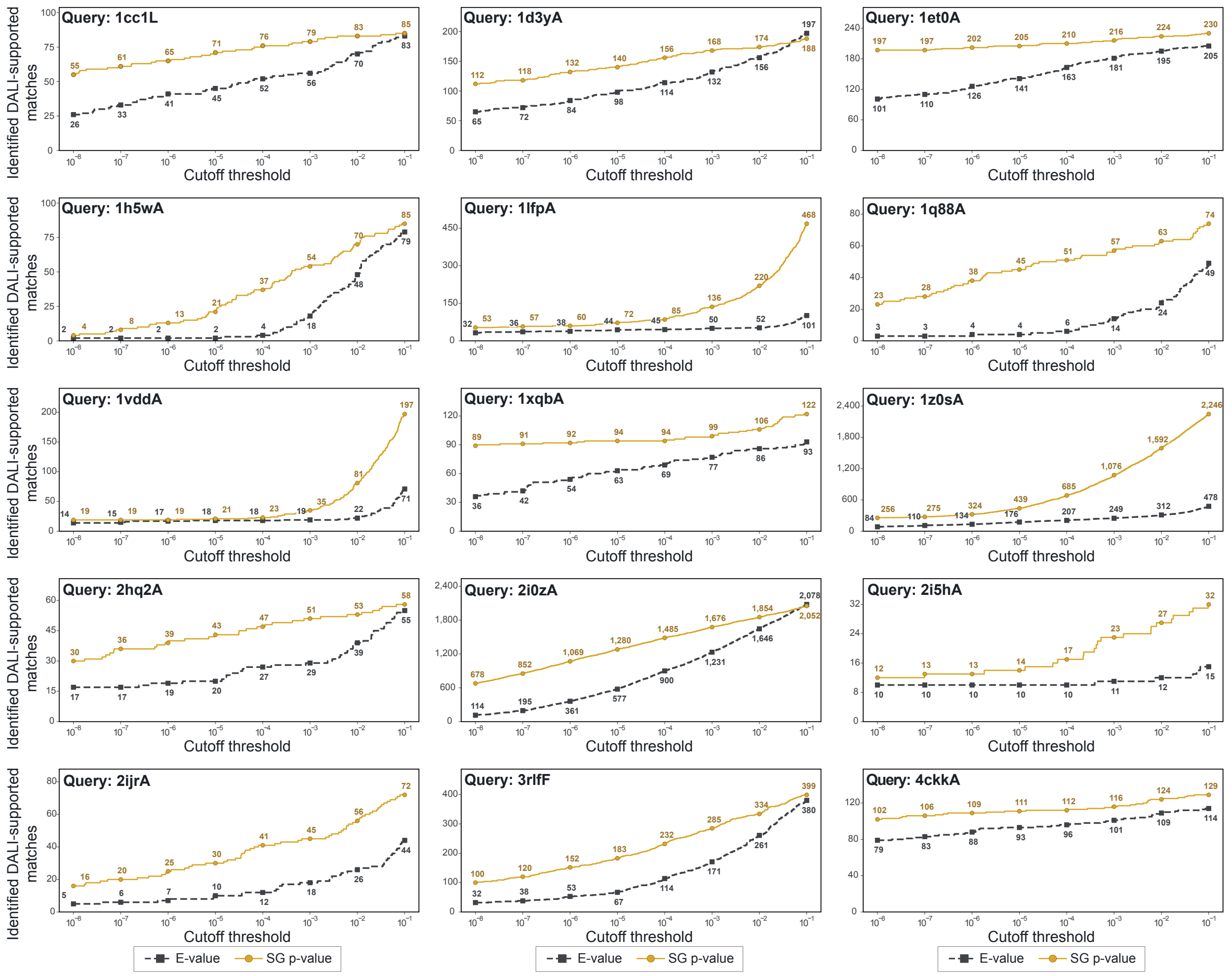

Figure S8: **Query-specific recovery of DALI-supported matches in AlphaFoldDB searches.** Each panel shows the number of DALI-supported matches recovered at different cutoffs for the indicated query, including `1h5wA` from the main analysis. Matches were considered structurally supported when their DALI scores met a reference threshold chosen separately for each query. Gray dashed lines with square markers show results using Foldseek native E-values, and yellow solid lines with circular markers show results using Struct-Caliper SG p-values. Numbers beside the markers give the counts at the eight indicated cutoffs, from $10^{-8}$ to $10^{-1}$. Horizontal axes are logarithmic, and vertical scales vary among queries.
